# An experimental approach to understanding elevation limits in a montane terrestrial salamander, *plethodon montanus*

**DOI:** 10.1101/131573

**Authors:** Nicholas M. Caruso, Jeremy F. Jacobs, Leslie J. Rissler

## Abstract

Understanding the abiotic and biotic factors that determine the limits to species’ range is an essential goal in ecology, biogeography, evolutionary biology, and conservation biology. Moreover, predictions of shifts in species’ distributions under future changes in climate can be improved through understanding the spatial variation in survival, growth, and reproduction. A long-standing hypothesis postulates that, for Northern Hemisphere species, abiotic factors like temperature limit northern and/or higher elevation extents, while biotic factors like competition limit the southern and/or lower elevation range edges; though amphibians may not follow this general trend. Therefore, we combined environmental suitability models and a reciprocal transplant experiment across an elevational gradient to explore the role of the abiotic environment on the range limits of a montane salamander (*Plethodon montanus*). We first determined suitability of the abiotic environment for *P. montanus*, under current (1960 – 2000) and future (2050) climate scenarios. Second, we collected juveniles from each of three elevations and transplanted them within mesocosms such that each origin population was represented within each transplant location and vice-versa. We found that environmental suitability in 2050 decreased throughout the range compared to current predictions, especially at lower elevations. Additionally, we found that individuals’ starting body condition and transplant location were important predictors of survival, growth, and reproduction condition; importantly, individuals transplanted to low elevation had lower survival and growth rates compared to those moved to mid or high elevations. Our study provides experimental support that the abiotic environment limits the lower elevation distribution of *P. montanus* and, unfortunately, our results also paint a possible bleak future for this species and likely other montane terrestrial plethodontids. The abiotic environment, which will become increasingly limited under future changes in climate, was found to have more influence on survival and growth than population identity.

## Introduction

Fundamentally, a species persists when the numbers of individuals entering the population (birth and immigration) are at least equal to those leaving (death and emigration) the population (Gaston 2009). However, the numbers and fitness of individuals across a species’ range vary due to spatial and temporal variation in abiotic (e.g., temperature, precipitation) or biotic (e.g., predation, food availability) factors. Limits to a species’ range often arise because of lowered resource availability or quality at range edges that can result in diminished condition or death of individuals (Hutchins 1947; Gaston 2003). For example, in the Southern Appalachians, warmer and drier conditions at lower elevations physiologically constrains the montane endemic salamander *Plethodon jordani* to higher elevations, while *P. teyahalee* is excluded from high elevations by the superior competitor, *P. jordani* (Gifford and Kozak 2012). Understanding how abiotic and biotic factors result in a population becoming a sink or extirpated is essential in determining the role of the environment on species’ range limits. As such it is both a fundamental question in ecology, biogeography, and evolutionary biology (Brown 1984; Gaston 2003; Case et al. 2005; Parmesan et al. 2005; Gaston 2009; Lee-Yaw 2016), and also critical for effective conservation and predictive modeling (Gaston 2003; Hampe and Petit 2005; Gaston 2009; Urban et al. 2016).

The North-South Hypothesis is a long-standing macroecological hypothesis (Darwin 1859; MacArthur 1972) which posits that abiotic conditions determine a species’ pole-ward or higher elevation range limit, but biotic conditions determine the equator-ward or lower elevation range limit (Dobzhansky 1950; MacArthur 1972; Brown et al. 1996; Gaston 2003; Parmesan et al. 2005; reviewed in Schemske et al. 2009; Hargreaves et al., 2014). For species in the Northern Hemisphere, this means that northern and/or higher elevation populations should be more constrained by abiotic factors like temperature, while southern and/or lower elevation populations should be more constrained by biotic factors like competition. However, several studies have shown that amphibians may not conform to this general trend (Hairston 1980; Nishikawa 1985; Gifford and Kozak 2012; Cunningham et al. 2016; Lyons et al. 2016). While studies examining range limits across different species are not uncommon (Schemske et al. 2009), there are fewer that examine what limits the different portions of a single species’ range (reviewed in Cahill et al. 2014; but see Cunningham et al. 2009).

Understanding the mechanisms responsible for range limits is also of concern for conservation biologists given the current rate of global climate change (Loarie et al. 2009). Mean global temperatures have increased by 0.6°C over the last century and are expected to increase 2 – 4°C more by 2100 (IPCC 2014). As a response to contemporary changes in climate, amphibians have shown shifts in breeding phenology (Beebee 1995; Reading 1998; Gibbs and Breisch 2001; Chadwick et al. 2006; Green 2016), geographic range limits (Pounds et al. 1999; Seimon et al. 2007), and body size (Reading 2007; Caruso et al. 2014). Environmental suitability models predict 50-100% reduction in suitable climate space for Southern Appalachians salamanders (Milanovich et al. 2010) – a major amphibian hotspot (Rissler and Smith 2010). Although potentially suitable future habitat may exist, many species lack the ability to disperse through the intervening lower suitability habitat in the face of climate change (Bernardo and Spotila 2006, Gifford and Kozak 2012; Lyons et al. 2016). Therefore, as global climates continue to shift, predicting species’ persistence will require an understanding of the relationship between climate and population vital rates (e.g., survival; Buckley et al. 2010; Urban et al. 2016) as well as decoupling local adaptation and plastic responses to climate (Parmesan 2006; Merilä and Hendry 2014; Urban et al. 2014).

Improvement to predictions of shifts in species’ distributions under future changes in climate can be accomplished through understanding how survival, growth, and reproduction vary spatially. Therefore, we combine environmental suitability models and a reciprocal transplant experiment across an elevational gradient to explore the role of the abiotic environment on the range limits of a montane salamander (*Plethodon montanus*). Using environmental suitability models we explore the variation in current and future suitability of the abiotic environment and compare models to identify how future ranges will be affected by changes in climate. Moreover, we used a reciprocal transplant experiment to investigate the role of origin population (i.e., where individuals were captured) and transplant location (i.e., where individuals were raised) on three relevant population responses: survival, growth, and maturation, to test the hypothesis that abiotic conditions limit the warmer range edge (i.e., lower elevations) of montane salamanders. Therefore, we asked the following questions, 1) how does environmental suitability vary across the current range of *P. montanus,* and how will suitability change (e.g., among elevations) under a future climate scenario? and 2) How do the origin location, transplant location, and initial body condition of individuals influence survival, growth rate, and maturation? Thus, if abiotic conditions limit the lower, warmer range edge, we expected to find lowest growth, survival, and maturation at the lower elevation sites and highest growth, survival, and maturation at the higher elevation sites. Moreover, if montane salamanders do best at their origin population we would expect to find increased survival, growth, and maturation for individuals at those locations compared to individuals transplanted to a non-origin location.

## Methods

### Environmental Suitability Models

We created predictive Ecological Niche Models (ENMs) for *P. montanus*, under current (1960 – 2000) and future (2050) climate scenarios, using program Maxent version 3.3.3k (Phillips et al. 2006). We obtained 262 unique high accuracy (at least four decimal points; ∼11 m resolution) geographic coordinates (latitude and longitude) using GBIF (<u>http://www.gbif.org/</u>) on 17 March 2016. We used the 11 bioclimatic variables that Rissler and Apodaca (2007) and Milanovich et al. (2010) used in distribution models of other plethodontid species (30 sec resolution; Hijmans et al. 2005) clipped to North America. We bootstrapped our ENMs by using 75% of the data for training and 25% of the data to test the models with 100 replicates. We used area under the curve (AUC) of the receiver operating characteristic (ROC) plots to evaluate model fit (i.e., values closer to 1 indicate a better fit; Swets 1988; Elith 2002), and determined if our current ENM models performed better than 262 random points with 999 replicates (Raes and ter Steege 2007). We used the default settings in Maxent for all other parameters and projected our current ENM model onto a future climate scenario (Hadley GEM2-ES model; RCP4.5 greenhouse gas scenario; 30 second resolution; Hijmans et al. 2005). For brevity, we report the average and standard deviation AUC, and data from the average and standard deviation of current and future ENM of bootstrapped models.

### Reciprocal Transplant Experiment

From 6 June – 1 October 2015, we conducted this transplant experiment in Pisgah National Forest, in which we collected salamanders who originated from low (∼1,000 m; 82.4258°W, 36.0328°N), mid (∼1,250 m; 82.1417°W, 36.1372°N), and high (∼1,450 m; 82.0917°W, 36.0928°N) elevations and reciprocally transplanted them to the same low, mid, and high elevation sites. Thus, each origin population was represented within each transplant location and vice-versa. Average annual temperatures varied predictably among these three elevations; the low site was the warmest (10.79°C), followed by the mid (9.59°C) and high elevation sites (8.78°C), whereas annual precipitation was lowest at the mid elevation site (1346.76mm) and higher and the low (1463.81mm) and high (1394.13mm) elevation sites (PRISM 2017). We established two replicate sites within each transplant elevation, with each replicate site containing 18 mesocosms (36 per elevation) for a total of 108 mesocosms. Replicate sites were chosen based on proximity to established mark-recapture sites, were approximately 200 – 400 m apart, and were on northwestern facing slopes with similar canopy coverage. Each mesocosm consisted of a single 53 x 43 x 30 cm polyethylene tub (Cunningham et al. 2009) that had the same number and size of holes drilled along the bottom and side for drainage. We filled each mesocosm with a layer of approximately 10 cm of soil, then a layer of approximately 2 cm of leaf litter (each gathered from the respective transplant site) and one 30 x 15 x 5 cm untreated pine cover board. The soil and leaf litter were collected and homogenized, separately, prior to adding equal amounts to each mesocosm in order to maintain consistency among mesocosms within each experimental site.

After establishing the mesocosms, we collected 36 juvenile salamanders that ranged from 30 – 45 mm SVL from each origin population. This size class was chosen because it represented the range of animals that could potentially reach reproductive maturity by the end of the experiment (N.M.C. *unpublished data*). Because the sex of juvenile salamanders is currently impossible to determine without dissection, we assumed a 1:1 sex ratio, which is consistent with museum specimens (N.M.C. *unpublished data*). Animals were kept in a cooler, maintained between 15 – 20°C, for approximately 36 hours before the start of the experiment. Immediately before beginning the experiment, we measured the snout-to-vent length (tip of the snout to posterior margin of the vent; SVL), tail length, and weighed each animal. All measurements were taken while the animal was secured in a new plastic bag to ensure consistent measurements and reduce probability of disease transmission from potentially contaminated equipment. We randomly assigned animals to a transplant elevation (low, mid, and high) and replicate site within transplant location (1 or 2). After adding a single salamander to each mesocosm, the mesocosm was covered with window screen and secured by both zip ties and waterproof caulk to prevent animal escape. Because we could not logistically collect all animals on a single day, the start date of the experiment varied from 6 June 2015 – 16 June 2015, and the end date of the experiment varied from 29 September 2015 – 1 October 2015 (106 – 115 days). At the end of the experiment, we thoroughly searched the leaf litter and soil of all mesocosms; salamanders were assumed dead if not found.

All animals were measured, euthanized (20% liquid Benzocaine), and dissected to determine sex and assess reproductive maturity. For males with pigmented testes, we assessed reproductive maturity by removing both testes and photographing them using a Leica M165C stereo microscope. All photos were taken on the same day, under identical lighting conditions, and using the same field of view. We used Leica Application Suite version 4.1 (Wetzlar, Germany) to determine the average area of both testes and ImageJ version 1.49 (Schneider et al. 2012) to determine mean pigmentation; we standardized testes area and pigmentation by the SVL at the end of the experiment, and we took the inverse of the mean standardized brightness of testes, such that darker testes would be scored as a higher number than lighter testes. Males with unpigmented testes were scored as 0 for standardized area and the inverse of mean testis brightness. Because testis area and inverse of pigmentation were correlated (0.65; t_22_=4.027; P < 0.001), we used testis pigmentation (scaled and centered) for further analyses. Although we designed our experiment to assess reproductive maturity for both sexes, no females were found with mature follicles; therefore, we did not assess female reproductive condition any further.

### Analyses

To determine if survival varied among origin, transplant, or initial body condition, we used a generalized linear mixed effect model with a binomial error distribution; our model contained two random intercepts, transplant location and replicate nested within transplant location. Similarly, we used linear mixed effects models, with the same random intercepts as above, to determine if growth rates (i.e., rate of change in SVL or mass) varied. To estimate body condition, we regressed log transformed mass against the total length (i.e., SVL + tail length) for all individuals (both beginning and end of the experiment) and extracted the residuals from the linear model; hereafter, we will refer to this as body condition index (BCI). A positive BCI indicates individuals with a greater mass for a given SVL, while a negative BCI indicates individuals with a lesser mass for a given SVL (e.g., Reading 2007; Băncilă et al. 2010). For both SVL and mass rates of change, we standardized these measures by first dividing them by the animal’s starting SVL and then dividing by the number of days that the animal was in the experiment. Lastly, we used a linear model, because of our smaller sample size, to determine if maturation (measured as testes pigmentation) varied among origin location, transplant location, or initial body condition. For our four response variables, we compared all 18 models using combinations of our three fixed effects (origin population, transplant location, and initial body condition) as well as interactions. Interactions were only included when the additive components of the interaction were present in the model. We were not able to fit the three-way interaction model for survival; therefore, it was not included. We fit all models using maximum likelihood and selected the best model based on lowest Akaike’s Information Criteria corrected for small sample size (AICc; Hurvich and Tsai 1989). We determined significance of model parameters of the top model using a likelihood ratio test. However, when competing models were within 2 AICc points of the top model (i.e., those that with “substantial support”; Burnham and Anderson 2002), we determined an average of model parameters, and the significance of parameters was determined as those with 95% confidence intervals that did not overlap zero; for brevity, we show visualizations for only significant results. For linear models (i.e., responses of SVL/mass change and male maturation), we graphically assessed models to ensure they met the assumptions of homoscedasticity and normality of residuals. All statistical analyses were performed in Program R version 3.3.1 (R Core Team 2016); we used the *dismo* package (Hijmans et al. 2016) to simulate random ENM models, the *lme4* package (Bates et al. 2015) for analyzing mixed effects models, the *Hmisc* package (Harrell Jr. et al. 2015) to determine binomial confidence intervals of apparent survival, and the *MuMIn* package (Barton 2016) to compare models by AICc, estimate average model parameters, and determine predictions from the model sets.

## Results

### ENM Modeling

Our ENM model provided an excellent fit (AUC = 0.992; SD = 0.001) to the locality data and was better than random (rAUC = 0.580 – 0.690). Within the current range of *P. montanus*, we found that the mean environmental suitability was 0.430 (SD = 0.044). Additionally, the occurrence data used to train and test our model had a mean ENM suitability of 0.518 (SD = 0.077). As expected, environmental suitability in 2050 decreased throughout the range (Fig. 1; Supplemental Fig 1), averaging 0.056 (SD = 0.034), while mean predicted ENM suitability was 0.112 (SD = 0.097) for the occurrence data used to train and test our models. Moreover, we found that the change in mean environmental suitability showed a linear or logistic trend associated with elevation while the difference in current and future suitability showed a quadratic or no relationship with both latitude and longitude (Fig. 1). Lastly, while current mean ENM suitability was similar for our focal sites (low = 0.570 [SD = 0.011); mid = 0.564 [SD = 0.007]; high = 0.572 [SD = 0.009]), predicted ENM suitability for 2050 is depreciated, especially at lower elevations (low = 0.075 [SD = 0.063); mid = 0.065 [SD = 0.054]; high = 0.202 [SD = 0.112]).

**Figure 1:**
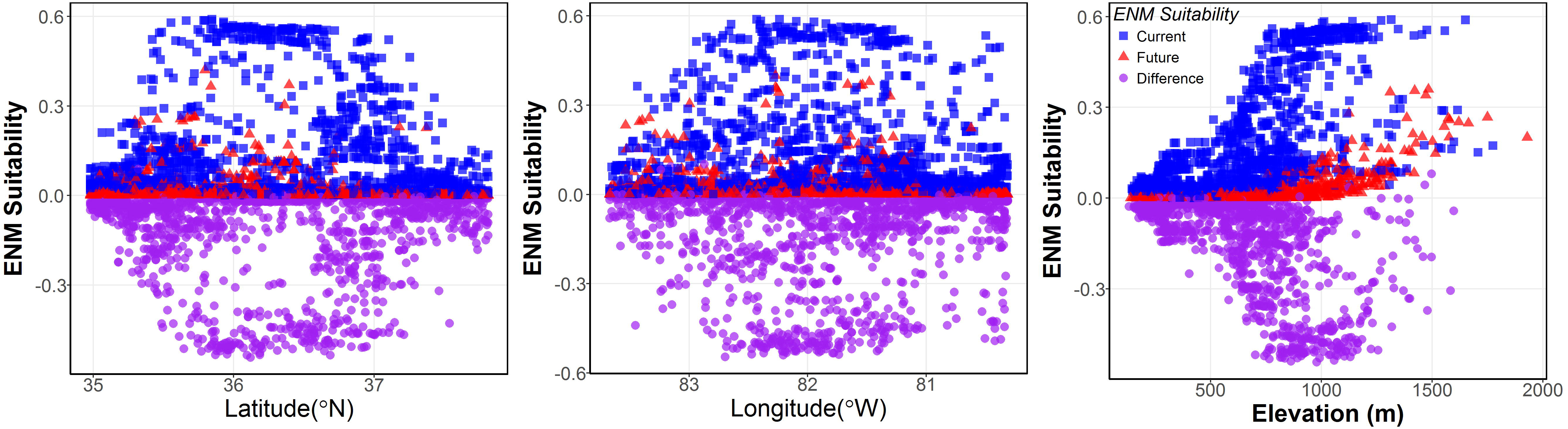
Relationship between ENM suitability and latitude, longitude, and elevation. Current (blue squares), future (red triangles) and the difference between future and current (purple circles) are shown.

### Reciprocal Transplant

We recovered 70 of the 108 salamanders at the end of the experiment (apparent survival = 65%; 95% CI = 55 – 73%). Apparent survival was lowest for individuals originating from higher elevations (56%; 95% CI = 40 – 71%) with increasing survival for animals from mid (64%; 95% CI = 48 – 78%) and low elevations (75%; 95% CI = 59 – 86%); however, apparent survival was highest for animals transplanted to higher and mid (72%; 95% CI = 56 – 84%) elevations compared to low elevation (50%; 95% CI = 34 – 66%). Individuals at the start of the experiment were generally smaller (mean SVL = 38.17 mm; 95% CI = 37.26 – 39.08 mm; mean Mass = 0.998 g; 95% CI = 0.933 – 1.062 g) but had a higher BCI (0.07; 95% CI = 0.03 – 0.11) compared to the end of the experiment (mean SVL = 42.64; 95% CI = 41.87 – 43.42 mm; mean Mass = 1.059; 95% CI = 1.002 – 1.115 g; mean BCI = -0.11; 95% CI =-0.13 – -0.08). Additionally, at the start of the experiment, SVL and mass were similar among origin populations and among origin populations and transplant locations. However, individuals transplanted to the low elevation gained less SVL and had more negative change in mass compared to those transplanted to mid and high elevations (Supplemental Figs. 2, 3). Regardless of origin population and transplant elevation individuals exhibited a negative change in BCI through the duration of the experiment (Supplemental Fig. 4). Lastly, most of the males (23/24; 96%) showed some degree of pigmentation and increase in size of their testes; the only male that did not show pigmented testes originated from, and was transplanted to, the low elevation site.

The most parsimonious predictors of survival were a set of five models containing the parameters of starting BCI, transplant location, origin population and the interaction between transplant locations and starting BCI (Table 1; Supplemental Table 1). The probability of survival was greatest for animals that started with a higher BCI, which is not unexpected. Interestingly, those salamanders with higher BCI who originated from low elevations and those that were transplanted to mid or high elevations had higher survival than other treatments (Fig. 2). For SVL, our top models included parameters of transplant location and starting BCI (Table 1; Supplemental Table 2). Animals that started off with a lower BCI had more positive rates of growth (SVL change), and this relationship was greatest for animals transplanted to mid elevations but lower for animals transplanted to high and low elevations (Fig. 3a). Though our model selection for mass change included more parameters, results were similar to the change in SVL (Table 1; Supplemental Table 3); animals that began the experiment with a lower BCI and were transplanted to mid and high elevations had a more positive rate of mass change (Fig. 3b). Lastly, we found that starting BCI significantly predicted (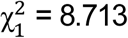; P < 0.001) maturation in males (Supplemental Table 4); males that started the experiment with a more positive BCI had darker testes (Fig. 4) and was not dependent on the environment or origin population.

**Figure 2:**
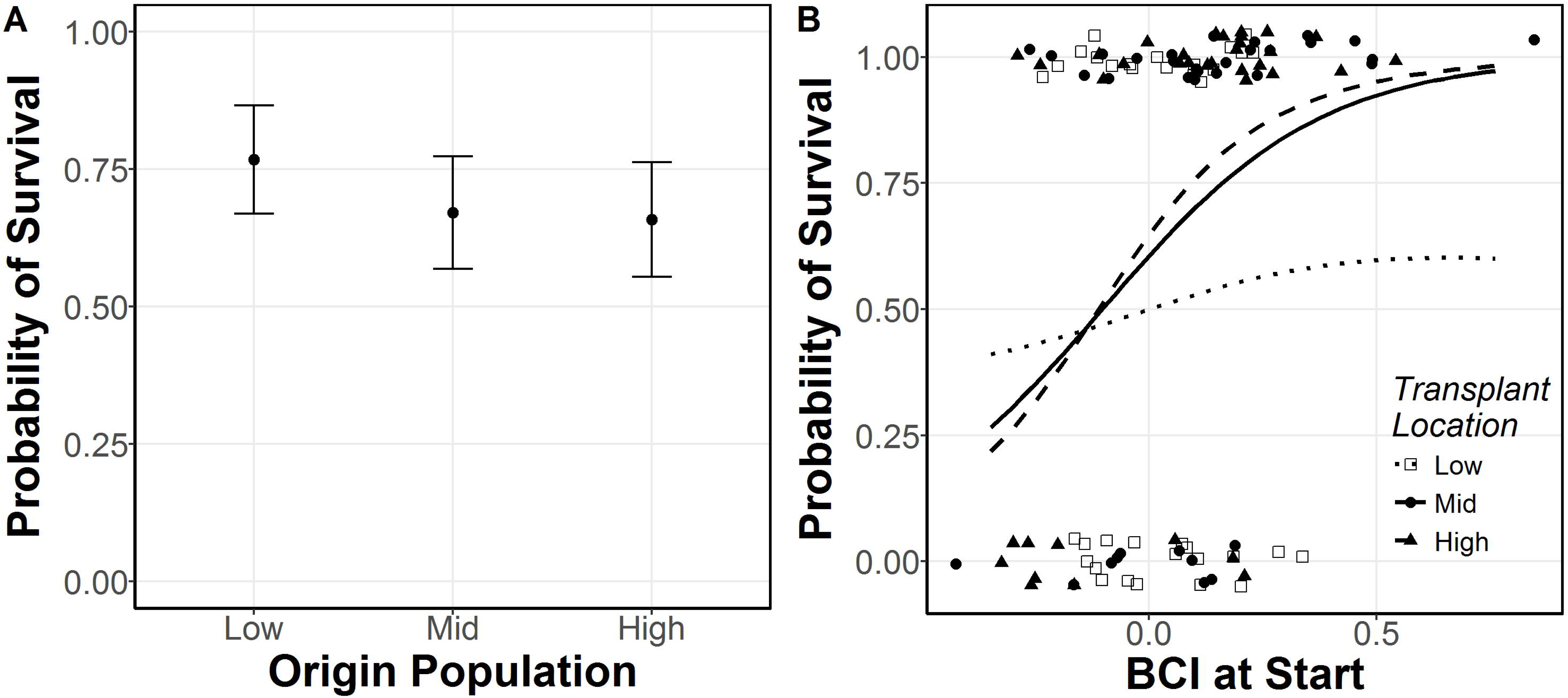
Relationship between survival of *P. montanus* and A) origin population, and B) transplant location and BCI. Error bars denote 95% CI, lines show predicted probability of survival and dots indicate data points for low (dotted line, open squares), mid (solid line, closed circles) and high elevations (dashed line, closed triangles).

**Figure 3:**
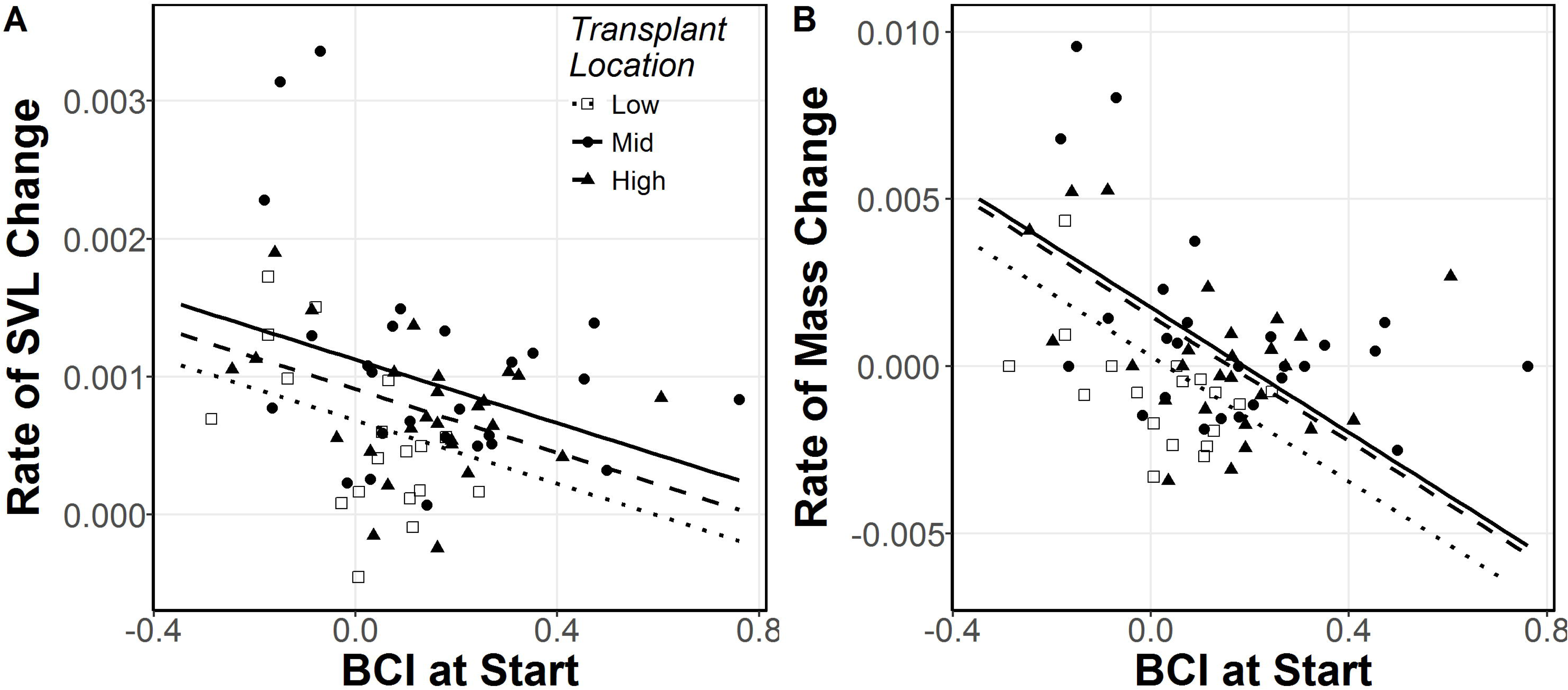
Influence of transplant location and BCI on rates A) of SVL and B) mass change. Lines show predicted fit, and dots indicate data points for low (dotted line, open squares), mid (solid line, closed circles) and high elevations (dashed line, closed triangles).

**Figure 4:**
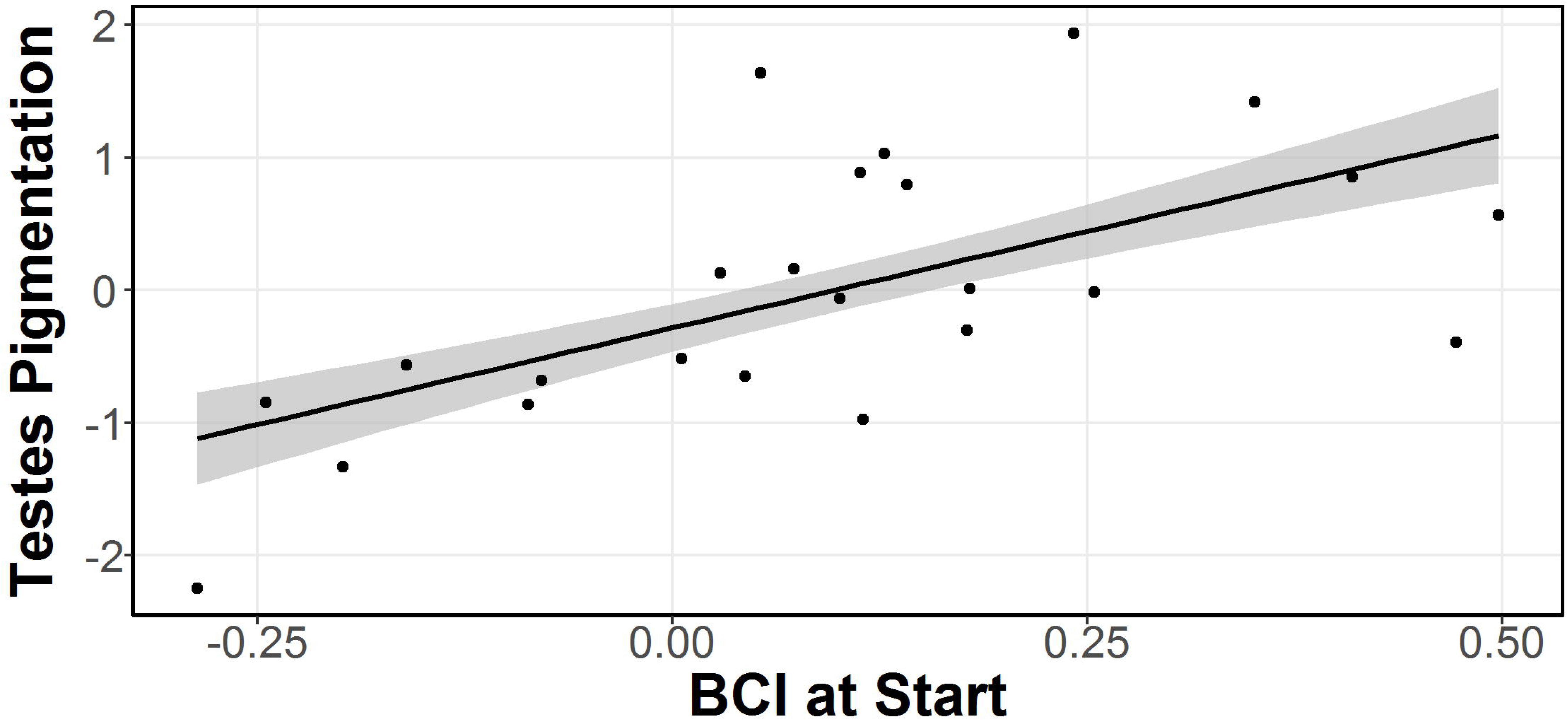
Reproductive condition (testes pigmentation) of *P. montanus* and BCI. Shaded ribbon denotes 95% CI of predicted fit (solid line).

**Table 1:**
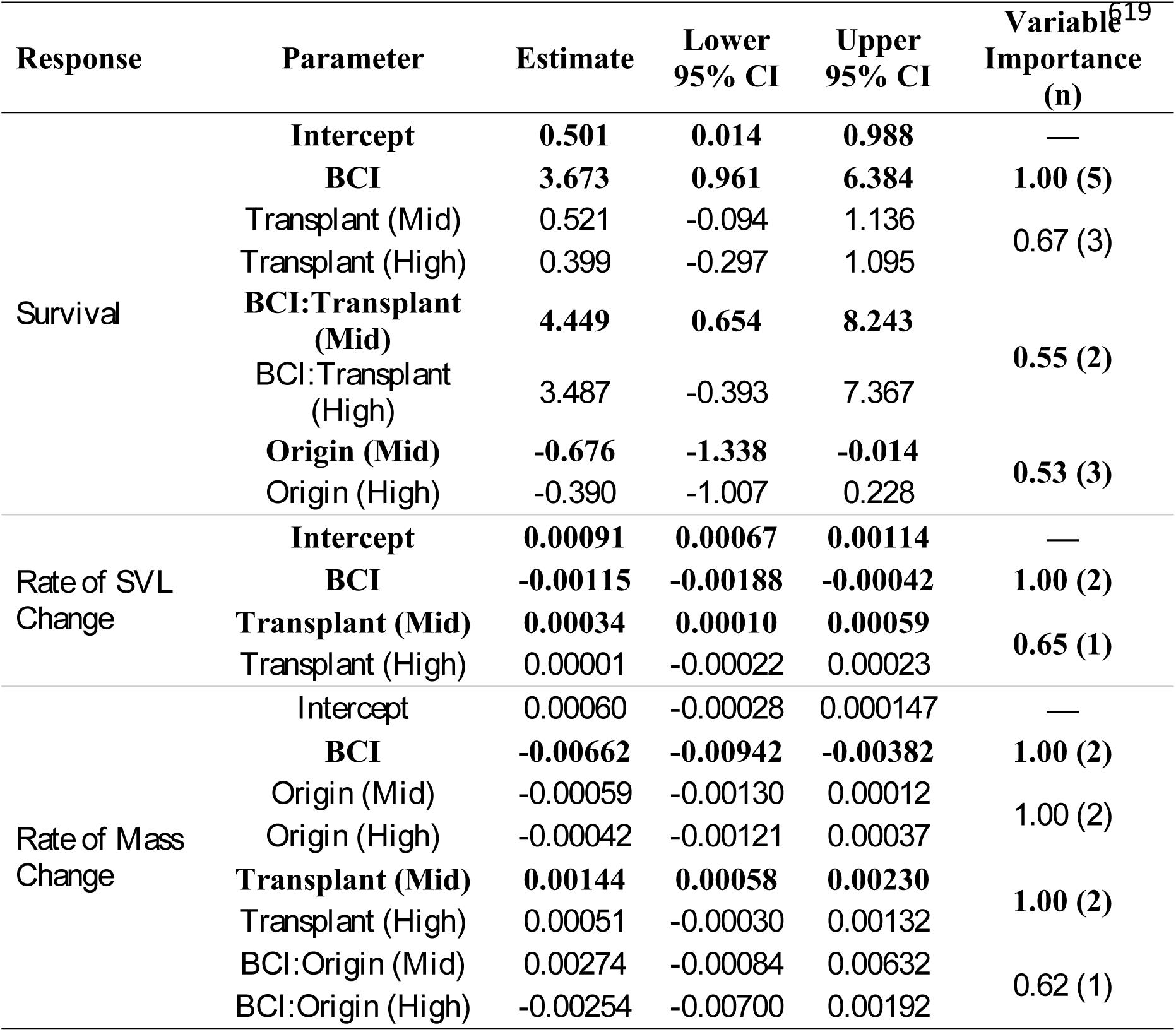
Estimates for the parameters included in the top model set (ΔAICc < 2) for each response (Supplemental Tables 1-4). Interactions are denoted by “:”, and bolded terms indicate significant variables (i.e., 95% CI do not overlap zero). Relative importance of each parameter is shown along with the number of models in the top model set that contain each variable (n).

| Response | Parameter | Estimate | Lower<br>95% CI | Upper<br>95% CI | Variable<sup>19</sup><br>Importance<br>(n) |
| --- | --- | --- | --- | --- | --- |
| Survival | **Intercept** | **0.501** | **0.014** | **0.988** | — |
|  | **BCI** | **3.673** | **0.961** | **6.384** | **1.00 (5)** |
|  | Transplant (Mid) | 0.521 | -0.094 | 1.136 | 0.67 (3) |
|  | Transplant (High) | 0.399 | -0.297 | 1.095 |  |
|  | **BCI:Transplant (Mid)** | **4.449** | **0.654** | **8.243** | **0.55 (2)** |
|  | BCI:Transplant (High) | 3.487 | -0.393 | 7.367 |  |
|  | **Origin (Mid)** | **-0.676** | **-1.338** | **-0.014** | **0.53 (3)** |
|  | Origin (High) | -0.390 | -1.007 | 0.228 |  |
| Rate of SVL Change | **Intercept** | **0.00091** | **0.00067** | **0.00114** | — |
|  | **BCI** | **-0.00115** | **-0.00188** | **-0.00042** | **1.00 (2)** |
|  | **Transplant (Mid)** | **0.00034** | **0.00010** | **0.00059** | **0.65 (1)** |
|  | Transplant (High) | 0.00001 | -0.00022 | 0.00023 |  |
| Rate of Mass Change | Intercept | 0.00060 | -0.00028 | 0.000147 | — |
|  | **BCI** | **-0.00662** | **-0.00942** | **-0.00382** | **1.00 (2)** |
|  | Origin (Mid) | -0.00059 | -0.00130 | 0.00012 | 1.00 (2) |
|  | Origin (High) | -0.00042 | -0.00121 | 0.00037 |  |
|  | **Transplant (Mid)** | **0.00144** | **0.00058** | **0.00230** | **1.00 (2)** |
|  | Transplant (High) | 0.00051 | -0.00030 | 0.00132 |  |
|  | BCI:Origin (Mid) | 0.00274 | -0.00084 | 0.00632 | 0.62 (1) |
|  | BCI:Origin (High) | -0.00254 | -0.00700 | 0.00192 |  |

## Discussion

Understanding how survival, growth, and reproduction vary spatially, leading to limits to species’ distributions, can inform population models and improve predictions of species’ range distributions under future changes in climate. Therefore, we determined the effect of origin population, transplant location, and initial BCI on juvenile salamander survival, growth, and reproductive condition. We found that individuals transplanted to mid and high elevations typically had higher survival and higher growth (SVL and mass) rates, while individuals that originated from low elevations had higher overall survival, irrespective of their transplant location (Figs. 2 – 4). Lastly, salamander body condition at the beginning of the summer is an important driver of survival, growth, and maturation; juveniles with a more positive BCI were more likely to survive, but had slower growth rates, and males had larger, darker testes (Figs. 2 – 4) at the end of the experiment.

### ENM Models and predicting future changes in suitability of sites and consequences to salamanders

Our current ENM models showed high and similar levels of suitability across our three transplant sites; however, ENM suitability for *P. montanus* decreased for forecasted 2050 climate, especially at lower elevations (Fig 1; Supplemental Fig. 1). Notably, mean predicted ENM suitability for 2050 was lower than 99.6% (261/262) of the *P. montanus* occurrence sites that we used to train and test our models. While our experiment was not conducted to explicitly test the role of environmental suitability per se on salamander growth and survival, which would require multiple transplant locations across many more sites across the environmental suitability landscape, our results do support predictions that lower elevations will become more limiting for montane salamanders as temperatures increase (Milanovich et al. 2010; Gifford and Kozak 2012; Lyons et al. 2016). Moreover, changes in climate do not affect species independently, and can result in shifts in the distribution of competitors, prey, or predators, which could alter interspecific interactions and potentially compound the negative effects of changes in climate on population growth (Blois et al. 2013; Liles et al. 2017).

### Initial Body Condition

Initial BCI was an important factor predicting observed trends in survival, growth, and reproductive condition in males; animals with higher BCI had higher survival rates, and males had higher reproductive condition. Our results for survival and reproductive condition agree with other studies that have examined the importance of body condition for individuals as well as populations (Wheeler et al. 2003; Karraker and Welsh Jr. 2006; Reading 2007; Janin et al. 2011). For example, Reading (2007) found that decreasing BCI in common toads, caused by warming climate, led to a decrease in survival and egg production. Our study shows that healthier individuals (i.e., those with higher body conditions) are afforded a greater probability of survival; however, for montane salamanders in low elevation habitats, this higher body condition may not be enough. As climate is expected to become less favorable for montane salamanders, especially at lower elevations, monitoring these low elevations populations is of increasing importance because at least 55% of montane plethodontids (lower elevation limit is greater than or equal to 1,000m) are threatened with extinction (IUCN 2016). Moreover, key life history characteristics such as body condition are relatively easy to collect and can provide an effective tool for identification of populations at risk (Janin et al. 2011).

Interestingly, animals that started the experiment with a greater BCI had lower rates of SVL changes and more negative changes in mass yet those same males had larger, darker testes indicative of greater reproductive condition (Sayler 1966; Peacock and Nussbaum 1973). This suggests that males with a greater BCI may put more energy into reproduction rather than growth during this life stage; this trend was strong for the males in our experiment (Supplemental Fig. 5). Tradeoffs between growth and reproduction in wild animals have been well-documented (e.g., Reznick 1983, 1985); for example, in *P. cinereus*, brooding females allocated less resources to growth compared to non-brooding females regardless of food availability (Yurewicz and Wilbur 2004).

### Origin Population

We found that where an individual originated was a significant predictor of survival; individuals from lower elevations had higher survival than individuals from mid and high populations (Fig. 2) irrespective of their transplant location. One of the limitations in our data was our inability to control for relatedness, maternal effects, or genotype-environment interactions (e.g., Via and Lande 1985; Sinervo 1990; Bernardo 1996; Bronikowski 2000), all of which could have added unknown sources of variation to our data. This was out of necessity as it would not have been feasible to assess reproductive condition in a natural setting for salamanders that take at least three years to mature (N.M.C. *unpublished data*). However, patterns of survival in our transplant experiment may suggest interacting effects of local adaption and phenotypic plasticity; individuals originating from low elevations were transplanted to elevations that represent either locally adapted conditions (i.e., low elevation) or better conditions (i.e., transplanted to higher elevations than origin). On the other hand, individuals who originated from mid and high elevations experienced locally adapted conditions or better conditions (when transplanted from mid elevation to high elevation only), or worse conditions (i.e., when transplanted to elevations lower than origin). This variation in survival may be explained, at least in part, by temperature, which decreased with increasing elevation (PRISM, 2017). Although low and mid elevations show similar ENM suitability, the scale at which these variables were measured (∼1 km) may not have been fine enough to adequately characterize these habitats. For example, Gifford and Kozak (2012) found that lower elevations, though they appeared to have identical habitats to higher elevations, contained microclimates that constrained *P. jordani*, a montane salamander. Future studies can refine hypotheses concerning local adaption and phenotypic plasticity in salamanders by splitting clutches or otherwise controlling for genetic factors; however, this technique is logistically challenging for many *Plethodon* species.

### Transplant Location

The abiotic environment, represented by transplant location in this experiment, was an important predictor for both growth and survival; salamanders that were transplanted to low elevations responded with the lowest survival and lowest growth (both mass and SVL) rates (Fig. 3). These results also conform to *in situ* observations; warmer summer temperatures were associated with reduced growth rates in *P. cinereus* (Muñoz et al. 2016) and lower elevations had lower survival than higher elevations in *P. montanus* (Caruso and Rissler 2017). Our results support the hypothesis that amphibian distributions are influenced at the southern or lower elevation range limits by abiotic variables (Buckley and Jetz 2007; Gifford and Kozak 2012; Cunningham et al. 2016; Lyons et al. 2016), although here we did not explicitly test the relative influence of biotic factors. It should be noted, however, that climatic barriers at the lower elevational limit are not universal for plethodontids and biotic variables may be more important. For example, *Desmognathus wrighti* is precluded from suitable lower elevations via predation by larger and more aquatic desmognathines (Organ 1961; Crespi et al. 2003), while *P. shendadoah* is outcompeted from non-talus slopes by *P. cinereus* (Jaeger 1970). Nonetheless we demonstrate here, that for the montane endemic *P. montanus*, the abiotic environment, specifically hotter and drier conditions (Fig. 1; Supplemental Fig. 1), likely limits its lower elevation distribution, similar to *P. jordani* (Gifford and Kozak 2012).

Though we did not modify or augment prey availability, we believe that prey availability was an unlikely source of variation in our results. First, all mesocosms had mesh screen and holes drilled along the side and bottom that were large enough to allow for smaller arthropods to enter the mesocosms. Second, local soil and leaf litter, which contained prey sources, were added to each mesocosm; thereby, minimizing variation among mesocosms within a transplant replicate. Lastly, we noted an abundance of prey items within the mesocosms when extracting the resident salamander at the end of the experiment, and during dissection to assess reproductive condition animals had prey items in their gut, indicating recent feeding (gut-passage time in *P. cinereus* is 1 – 2 weeks; Merchant 1970; Gabor and Jaeger 1995). While we do not suppose that prey populations affected our experiment, future shifts in salamander abundance could have big consequences for ecosystem function. Due to their large numbers in forest ecosystems (Burton and Likens 1975; Milanovich and Peterman 2016), salamanders can exhibit strong top-down effects on invertebrate populations; their presence is associated with reduced leaf-litter decomposition and overall carbon retention (Wyman 1998; Rooney et al. 2000; Walton et al. 2006; Best and Welsh 2014; but see Walton and Steckler 2005; Homyack et al. 2010; reviewed in Walton 2013). Therefore, declines of salamander populations may have a negative feedback, in which the loss of salamander biomass reduces forest carbon sequestration, potentially accelerating anthropogenic climate change, resulting in further reductions in areas of suitable climate for salamander persistence.

In conclusion, we first provide experimental support for the hypothesis that the abiotic environment constrains the lower elevation limits of *Plethodon montanus*, which is consistent with predictions for amphibians, and more specifically, montane salamanders (e.g., Gifford and Kozak 2011; Cunningham et al., 2016; Lyons et al. 2016. Importantly, by using a reciprocal transplant experiment, we were able to test the relative influence of origin population and transplant location simultaneously; our results (i.e., AICc-selected variables and variable importance; Table 1; Supplemental Tables 1-3), suggest that the abiotic environment (transplant location) has more influence on survival and growth than population identity (origin population). Warmer and/or drier conditions can result in reduced surface activity, increased metabolism, increased water loss, as well as reductions in growth and survival in plethodontids (Caruso et al. 2014; Riddell and Sears 2015; Catenazzi 2016; Connette et al. 2015; Muñoz et al. 2017; Caruso and Rissler 2017). Continued trends towards warmer and drier climates in the Appalachian region will likely lead to reductions in population growth unless compensated by an increase in immigration or reproduction (Tavecchia et al. 2016; Gaston 2009). These data are important not just because they add to the growing body of literature seeking to understand what determines the limits of a species’ range (Gaston 2003), but they further suggest a worrisome forecast for montane salamanders under predicted future climate changes.

## Acknowledgements

Thanks to P. Scott and S. Duncan for assistance setting up the mesocosms and to C. Staudhammer, G. Starr, D. Adams, P. Scott, S. Wiesner, S. Kunwor, and S. George for comments that improved this manuscript. This research was funded through Graduate Research Fellowship, E.O. Wilson Fellowship, and the Herpetologists’ League E.E. Williams Research Grant awarded to NMC. We had NC state and Pisgah N.F. permits to conduct this research. All animal work was conducted as outlined by national guidelines (University of Alabama IACUC approval 15-02-0098). The authors declare no conflict of interest.

